# Genetic mapping in Diversity Outbred mice identifies a *Trpa1* variant influencing late phase formalin response

**DOI:** 10.1101/362855

**Authors:** Jill M. Recla, Jason A. Bubier, Daniel M. Gatti, Jennifer L. Ryan, Katie H. Long, Raymond F. Robledo, Nicole Glidden, Guoqiang Hou, Gary A. Churchill, Richard S. Maser, Zhong-wei Zhang, Erin E. Young, Elissa J. Chesler, Carol J. Bult

**Affiliations:** The Jackson Laboratory 600 Main Street, Bar Harbor, ME 04609, USA; IGERT Program in Functional Genomics Graduate School of Biomedical Sciences and Engineering The University of Maine Orono, ME 04469, USA; Department of Genetics and Genome Sciences UCONN Health 400 Farmington Avenue Farmington, CT 06030-6403, USA; School of Nursing University of Connecticut 231 Glenbrook Rd, Unit 4026 Storrs, CT 06269-4026, USA; Institute for Systems Genomics University of Connecticut Storrs, CT 06269-4026, USA

**Keywords:** pain genetics, formalin, chronic pain, chemical nociception, Diversity Outbred mice, genetic linkage mapping

## Abstract

Identification of genetic variants that influence susceptibility to chronic pain is key to identifying molecular mechanisms and targets for effective and safe therapeutic alternatives to opioids. To identify genes and variants associated with chronic pain, we measured late phase response to formalin injection in 275 male and female Diversity Outbred (DO) mice genotyped for over 70 thousand SNPs. One quantitative trait locus (QTL) reached genome-wide significance on chromosome 1 with a support interval of 3.1 Mb. This locus, *Nociq4* (nociceptive sensitivity inflammatory QTL 4; MGI:5661503), harbors the well-known pain gene *Trpa1* (transient receptor potential cation channel, subfamily A, member 1). *Trpa1* is a cation channel known to play an important role in acute and chronic pain in both humans and mice. Analysis of DO founder strain allele effects revealed a significant effect of the CAST/EiJ allele at *Trpa1*, with CAST/EiJ carrier mice showing an early, but not late, response to formalin relative to carriers of the seven other inbred founder alleles (A/J, C57BL/6J, 129S1/SvImJ, NOD/ShiLtJ, NZO/HlLtJ, PWK/PhJ, and WSB/EiJ). We characterized possible functional consequences of sequence variants in *Trpa1* by assessing channel conductance, *Trpa1/Trpv1* interactions, and isoform expression. The phenotypic differences observed in CAST/EiJ relative to C57BL/6J carriers were best explained by *Trpa1* isoform expression differences, implicating a splice junction variant as the causal functional variant. This study demonstrates the utility of advanced, high-precision genetic mapping populations in resolving specific molecular mechanisms of variation in pain sensitivity.

## INTRODUCTION

Chronic pain is a maladaptive condition in which the sensation of pain persists in the absence of an eliciting stimulus. It is estimated to affect up to 30% of the world’s population [26]. With a reported trait heritability of 16-50% in humans [34; 60], the onset and continuation of chronic pain is influenced heavily by genetic background. Pain-related genetic variants identified to date influence variation in neurotransmitters and their receptors, growth factors, inflammatory cytokines, and myriad other neuromodulators [28; 88]. Although several highly-penetrant human genetic variants are known to underlie rare familial monogenic pain conditions [28; 43; 65; 68; 84; 89], the genetic landscape of common chronic pain conditions suggests minor contributions from a large number of single nucleotide polymorphisms (SNPs) representing diverse functional pathways [88; 89].

The laboratory mouse has proven to be a useful discovery platform for the genetic study of human chronic pain; findings from several mouse studies have been corroborated in humans [54-56; 61; 67; 77]. Low allelic variation, genetic recombination density and resulting lack of mapping precision, however, limit the utility of conventional mapping strategies using the laboratory mouse for discovery of new genes and variants related to pain phenotypes. The Diversity Outbred (DO) stock [21] is a mouse population derived from a set of eight genetically diverse parental strains (A/J, C57BL/6J, 129S1/SvImJ, NOD/ShiLtJ, NZO/HlLtJ, CAST/EiJ, PWK/PhJ, and WSB/EiJ) that has increased heterozygosity and allelic diversity compared to conventional mapping populations. DO mice are produced by the repeated random outcrossing of non-siblings originally from the Oak Ridge National Laboratory (ORNL) Collaborative Cross (CC) colony [20], a genetically defined panel of recombinant inbred lines. The DO mouse population captures a large set of natural allelic variants derived from a common set of eight founder strains, providing multitudinous combinations of segregating alleles in virtually all genetic loci in the mouse genome [76]. The high genetic diversity and precision afforded by the DO makes it an ideal resource for tractable identification of novel genes and variants governing chronic pain. We published the first application of DO mice and genetic linkage mapping to the study of pain genetics in 2014 [66], where we identified a novel role for a single, protein-coding candidate pain gene, *Hydin* (HYDIN, axonemal central pair apparatus protein; MGI:2389007), postulated to influence thermal pain response via a previously unreported ciliary mechanism in the choroid plexus–cerebrospinal fluid system.

Here, we build upon our previous work by examining a chronic pain model, the late phase response to formalin injection. We used updated statistical algorithms for genetic linkage mapping and SNP association mapping in CC and DO mice [76] to map genetic loci involved in early and late phase formalin response, and identify precise allelic variants in the DO population that could be responsible for variation in pain response. We then experimentally evaluated the causal mechanisms attributable to specific variants in the QTL to determine which was responsible for variation in the pain response.

## MATERIALS AND METHODS

### Diversity Outbred Mice

Male and female DO mice (n =300; J:DO, JAX stock number 009376) from generation 8 (G8) of outcrossing were obtained from The Jackson Laboratory at 11 weeks of age. Mice were transferred from the breeding facility directly to an adjoining housing facility via wheeled cart and were acclimated to the vivarium for at least 2 weeks prior to testing at 13–17 weeks of age. Mice were housed in duplex polycarbonate cages with a Shepherd Shack® on ventilated racks providing 99.997% HEPA filtered air to each cage in a climate-controlled room under a standard 12:12 light–dark cycle (lights on at 0600 h). Pine cob bedding was changed weekly and mice were provided ad-libitum access to food (NIH31 5K52 chow, LabDiet/PMI Nutrition, St. Louis, MO, USA) and acidified water. Female mice were group-housed with a cage density of 4-5 individuals per cage. Male mice were single housed, as earlier studies indicate a propensity toward aggressive behavior in group-housed DO males [47; 66]. All procedures and protocols were approved by The Jackson Laboratory Animal Care and Use Committee (Bult AUS #01011) and were conducted in compliance with the National Institutes of Health Guidelines for the Care and Use of Laboratory Animals.

### Experimental Design

A total of 288, 13-17 week old DO mice (147 female, 141 male) were phenotyped using the formalin assay of nociception. Twelve of the original 300 mice were excluded from analysis due to bite wounds or congenital abnormalities, including hind leg splay and cranial malformation. Mice were randomly assigned to testing groups, such that an equal number of male and female mice were tested each day (n = ∼16 per sex). Two groups were tested per day during the animals’ resting phase. The morning group tested between 09:00 and 10:00 h, the afternoon group between 11:00 and 12:00 h. A single experimenter performed all of the injections, and the testing room was vacated for the duration of the assay.

### Formalin Assay of Nociception

Mice were transported to the testing room (maintained at 24-25°C) and left undisturbed in their polycarbonate cages to habituate for 1 hour under ambient light before beginning the experiment. Twenty microliters of 2.5% formalin solution was injected subcutaneously into the plantar surface of the left hind paw using a 0.3 cc micro-syringe (Hamilton) with a 30-gauge needle. Injected mice were individually placed into 4-quadrant plastic observation chambers (4”w x 4”l x 5”h), located on a flat, glass surface to allow clear observation of the injected paw. Administration of 2.5% formalin was sufficient to produce the desired biphasic response while increasing test sensitivity and reducing animal suffering compared to the most commonly used 5% formalin solution.

Noldus Observer 2.1 (Noldus Information Technology, Wageningen, The Netherlands) was used to record video with one camera per observation chamber mounted below the glass surface. We recorded the time spent licking or biting the formalin-injected paw in 1-min intervals up to 60 min beginning immediately after formalin injection after the time sampling method [2]. Stationary video cameras were used to record the behavioral responses. Video observations were binned into 60 sec time points and scored manually by a single trained investigator. A time point was assigned a score of 1 if the mouse was observed licking or biting the injected paw, 0 otherwise. Scores were summed across time points for the acute (0-10 mins) and chronic (11-60 mins) response phases, giving each mouse an acute pain score of 0-10 and a total chronic pain response score of 0-50. Longer time spent licking or biting the injected paw during the late response phase was taken to imply increased susceptibility to chronic pain.

### Genotyping

Genotyping was performed on all 288 DO samples. DNA was prepared from tail tips and genotyped using the second generation Mouse Universal Genotyping Array (MegaMUGA) performed by the GeneSeek service (http://www.neogen.com/GeneSeek; Lincoln, NE, USA). Built on the Illumina Infinium platform (San Diego, CA, USA), the MegaMUGA contains 77.8K SNP markers distributed throughout the mouse genome with an average spacing of 33 Kb [23; 81]. SNPs were selected to be representative of the diversity in the founding strains of the CC and DO – A/J, C57BL/6J, 129S1/SvImJ, NOD/ShiLtJ, NZO/HlLtJ, CAST/EiJ, PWK/PhJ, and WSB/EiJ [58].

### QTL mapping

QTL mapping was performed in 275 DO mice (147 female, 128 male). Of the 288 phenotyped animals, three mice were excluded due to missing genotype calls and ten were excluded due to video recording error. Mapping was carried out as described by Gatti et al. [29]. All phenotype and genotype data have been made publicly available through the QTL archive at the Mouse Phenome Database under study name Recla2 (MPD, http://phenome.jax.org/) [16].

#### Additive haplotype model

The additive model assumes that each copy of the founder alleles contributes a unit of trait variation; there are no dominance effects in this model. Founder haplotypes were reconstructed using a Hidden Markov Model (HMM) that produced a matrix of 36 genotype probabilities for each sample at each SNP. Genotype probabilities at each SNP were then collapsed to an eight-state allele dosage matrix by summing the probabilities contributed by each founder. Phenotypic data were normalized by square root transformation of total chronic pain response score prior to linkage mapping analysis to satisfy model assumptions. Mapping was performed using QTLRel software (http://www.palmerlab.org/software) [19]. A mixed model was fit with sex and AM/PM group as additive covariates and a random effect was included to account for kinship. Regression coefficients for additive effects of founder haplotypes were estimated at each genomic location. Significance thresholds were obtained by performing 1000 permutations of the genome scans with phenotype data being shuffled among individuals and 2-LOD support intervals from the linear model were determined for significant (*p* ≤ 0.05), suggestive (*p* ≤ 0.10), and trending (*p* ≤ 0.63) QTL peaks.

#### Additive SNP model

The SNP-based additive model is widely used in human association mapping [17]. Mapping at the two-state SNP level increases power and precision by assessing the effects at individual variants, and has the potential benefit for evaluating dominance effects with one additional degree of freedom [29]. To implement the additive SNP model, we computed a probabilistic imputation of the genotype at every known SNP locus genome-wide. We then fit an additive SNP model by regressing the square root of the total chronic pain response score on the imputed DO genotypes. Mapping was performed using QTLRel software (http://www.palmerlab.org/software) [19]. To gain computational efficiency, we assigned the diplotype state probability between each adjacent pair of genotyped markers to the average of the flanking diplotype state probabilities. Any sets of SNPs in the interval with identical strain distribution patterns among the eight founders were assigned identical values and we computed the regression once for each set of identical SNPs in an interval. Significant SNPs were determined to lie within a 1-LOD support interval from the maximum LOD.

### Candidate gene analysis

To assess the plausibility of candidate genes in the *Nociq4* region we compiled functional, phenotypic, and expression annotations from a variety of databases (Supplemental Table S1) using methods for candidate gene prioritization we developed previously [66]. First, we identified all protein-coding and functional RNA genes within the *Nociq4* region (chr1:11.95..15.07 Mb) using the unified mouse gene catalog from the Mouse Genome Informatics (MGI) database (http://www.informatics.jax.org/marker/) [15]. Second, for each genome feature in the region, we compiled functional, phenotypic, and expression annotations from the informatics resources in Table S1 as follows: gene expression annotations were collected from the Allen Brain Atlas (ABA) [46], EBI Expression Atlas (EEA) [63], Gene Expression Omnibus (GEO) [10], and the Gene eXpression Database (GXD) [71] through MGI [15]; functional InterPro protein domain [27], Mammalian Phenotype (MP) [70], and Gene Ontology (GO) annotations [6; 78] were obtained through MGI; pain-related phenotype data from pain gene knock-out models were collected from MGI and PainGenesdb [42]. Finally, SNP locations from the Sanger Mouse Genomes Project version 5 (REL-1505) [37] were used to identify SNPs in the additive SNP model significantly associated with *Nociq4*. Gene annotations from Ensembl annotation version 75 [85] were used to computationally plot the candidate genes underlying each SNP.

To identify plausible genetic and functional candidate genes in a computationally predictive manner, sets of genes were created in GeneWeaver [7] using the MGI and Ensembl gene lists generated above. The GeneSet Graph tool was used to intersect the genes in the QTL interval with those that have SNPs specific to the CAST/EiJ strain. In addition, we intersected a set of genes derived from RNA-Seq data (GEO GSM2743739) to identify genes that are expressed in the dorsal root ganglia (DRG). Finally, we interested a set of 889 mouse genes associated with the Mammalian Phenotype Ontology term (MP:0002067) “abnormal sensorial capabilities/reflexes/nociception.” The resulting GeneSet graph produced by GeneWeaver predicts the most likely candidate gene given the conditions described above.

### Phenotypic contributions of DO founder strains

We examined the relationship between allelic variation and phenotypic response at *Nociq4* by first computationally sorting the original DO mapping population into groups based on the parental allele at the *Nociq4* peak (Chr1:14.25 Mb; n=8, 1 group per DO founder strain). We then calculated the mean allelic response for each DO founder strain by averaging the pain response scores per group at each 1 min time point over the 60 min formalin testing period. Results were calculated and plotted over time in R software environment 3.0.2.

### Trpa1 SNP analysis

A major benefit of the DO in mapping studies is the ability to precisely associate observable phenotypic variation with specific underlying genetic variants. We identified putative causal variants unique to CAST/EiJ in *Trpa1* by examining the imputed DO genotype data used to fit the additive SNP model at *Nociq4*. We selected SNPs with LOD scores greater than the maximum LOD score (5.71) minus one and intersected them with the exons and untranslated regions of *Trpa1*, obtained and processed computationally from NCBI dbSNP Build 150 [1; 69]. We further subset these SNPs by selecting missense, splice site, or other regulatory variants likely to produce a functional consequence (Sanger Mouse Genomes Project version 5; REL-1505 [37]).

### Electrophysiological evaluation of ankyrin domain variant

CAST/EiJ variant rs32035600 induces a Valine to Isoleucine codon shift (Val115Ile) in the *Trpa1* ankyrin repeat domain (ARD), a region of the folded protein known to influence the electrophysiological properties of the channel [32; 44; 83], and which is involved in the aggregation of *Trpa1* and *Trpv1* receptors. We evaluated the electrophysiological consequence of CAST/EiJ variant rs32035600 on *Trpa1* channel conductance using whole-cell patch clamp recording in HEK293T cells.

### Cell line mutation

We obtained clone EX-Mm17807-M03 (Genecopoeia) for *Trpa1* ORF driven by a CMV promoter and a C-terminal GFP tag. Using site directed mutagenesis in *E.coli*, mouse SNP rs32035600 was mutated from the C57BL/6J (G) to the CAST/EiJ (A) variant. Both variants were transfected into human embryonic kidney cells (HEK293T cells). HEK293T cells were grown under standard conditions in DMEM with 10% FBS, Glutamax, and penicillin/streptomycin, and were transfected with expression plasmids encoding GFP-tagged *Trpa1* carrying the two variants of interest. Cells were transfected using 500 ng of plasmid DNA and 1.5 ul of Lipofectamine 3000 transfection reagent, according to the manufacturer’s protocols.

### Patch-clamp analysis

Forty-eight to 72 h after transfection, HEK293T cells grown on glass coverslips were transferred to a submersion chamber where they were continuously perfused with extracellular recording solution containing (in mM): 124 NaCl, 3.0 KCl, 1.5 CaCl_2_, 1.3 MgCl_2_, 1.0 NaH_2_PO_4_, 26 NaHCO_3_, and 20 glucose, saturated with 95% O_2_ and 5% CO_2_ at room temperature (21-23 °C). Cells were viewed with a 40x objective (N.A. 0.8, water immersion) and transfected cells were identified using GFP epifluorescence. Whole-cell patch clamp recordings were performed with a Multiclamp 700B amplifier (Molecular Devices, Sunnyvale, CA). The pipette solution contained (in mM): 130 CsCl, 4 ATP-Mg, 0.3 GTP-Na, 0.5 EGTA, and 10 HEPES (pH 7.2, 270-280 mOsm with sucrose). The series resistance, usually between 7-12 MΩ, was continuously monitored but not compensated. Data were discarded when series resistance changed by more than 25% during the experiment. Mustard oil (allyl isothiocyanate, Sigma-Aldrich, #377430) was diluted in the extracellular solution to the concentration of 200 μM and applied through a buffer pipette placed 40-50 μm away from the recorded cell. The buffer pipette had a tip diameter of 2 μm and the pressure pulses were 10 s long at 20 psi. Experiments were conducted using AxoGraph X (AxoGraph Scientific, Sydney, Australia). Data were filtered at 2 kHz and digitized at 8 kHz. Data analysis was performed using AxoGraph X.

### *Evaluation of ankyrin binding interactions by Co-Immuno-Precipitation of* Trpa1-Trpv1

The Val115Ile codon shift induced by CAST/EiJ variant rs32035600 may affect the ability of *Trpa1* to bind and co-localize with its functional partner *Trpv1* (MGI:1341787; transient receptor potential cation channel, subfamily V, member 1) [72]. To test this, we performed Co-Immuno-Precipitation of the *Trpa1-Trpv1* complex in mouse DRG from male and female formalin and saline treated CAST/EiJ and C57BL/6J mice. Tissue extracts were prepared using Lysis Buffer with Protease Inhibitors (LBPI; 300 μL/sample). Samples were then homogenized with a mortar/pestle, incubated at 4**°** C with gentle agitation for one hour followed by centrifugation (Eppendorf 5417C) at 20000 x g for 20 minutes at 4**°** C to remove cell debris. The supernatant/tissue extract was then transferred to a fresh microcentrifuge tube. *Trpv1* antibody (1 μg total; Abcam, Cat#ab6166) was added to each normalized cell lysate sample (100μg/μL). Fresh LBPI was added to each sample to reach a final volume of 500 μL. Samples and antibody incubated overnight at 4**°** C with gentle agitation. Protein A/G Sepharose bead slurry (75 μl/sample; Santa Cruz Cat# sc-2003) was added to each tube for overnight incubation at 4**°** C with gentle agitation. Agarose beads were collected by (8000 x g) centrifugation and were washed (x3) with 1 mL 1X Wash Buffer. 2X SDS/PAGE loading buffer (30 μL) was added to the beads and samples were boiled for 5 minutes to elute the complex. Eluent was loaded directly into single wells in a 4-15% acrylamide gel (Bio-Rad) and run for 44 minutes at 200 volts followed by transfer to a nitrocellulose membrane for 1 hour at 100 volts. The nitrocellulose blot was probed for the presence of *Trpa1* (EMD Millipore, Cat# ABN1009; 1:1000) and β*-actin* (Actin Novus Biologicals, Cat# NBP254690; 1:5000). The blot was also probed for the presence of *Trpv1* (Alomone Labs, Cat# ACC-030; 1:1000) and β*-actin* (Actin Novus Biologicals, Cat# NBP254690; 1:5000). ImageJ (U. S. National Institutes of Health, Bethesda, MD, USA) was used to quantify the band intensities. This experiment was replicated a second time. In each replicate the ratio of β*-actin*-normalized *Trpa1*/*Trpv1* was obtained per sample.

### *qRT-PCR expression analysis of* Trpa1

CAST/EiJ SNP rs239908314 is a predicted splice region variant located in *Trpa1* intron 21. Exon 20 is skipped in mouse splice variant *Trpa1*b, implicating rs239908314 in *Trpa1* isoform transcript control. The full-length *Trpa1* transcript, designated *Trpa1*a, interacts physically with *Trpa1b* to enhance *Trpa1*a expression on the plasma membrane, significantly increasing *Trpa1* agonist responses [86]. We explored *Trpa1* isoform transcript expression in DRG from pain-sensitive (C57BL/6J; n=7, 3 male) and pain-resistant (CAST/EiJ; n=8, 4 male) mice. DRG were dissected and stored in RNAlater. RNA was extracted using TRIzol reagent, quality assessed using Agilent Bioanalyzer Nano Chips. 200ng total RNA was converted to cDNA using random-decamers. TaqMan assays were used to measure abundance of *Trpa1* exon 13-14 (TaqMan Assay ID Mm00625257_ml) common to *Trpa1*a and *Trpa1*b and exons 19-20 (Mm01227443_ml) and exons 20-21 (Mm00625257_ml) both specific for *Trpa1*a. The threshold cycle (Ct) for each probe was determined using the ViiA 7 software (Thermo Fisher Scientific, Waltham, MA, USA). The data were further analyzed using the ΔΔCt method, normalized to *Gapdh* (Mm99999915_g1), and plotted in Fig 8. qRT-PCR data are archived in MPD (http://phenome.jax.org/) [16].

## Results

### Genetic Linkage Mapping and SNP Association Mapping Identify a Single QTL Peak on Chromosome 1

Genetic linkage mapping and SNP association mapping identified one single QTL peak of genome-wide significance for late phase response to formalin injection located on chromosome 1, which we have named nociceptive sensitivity inflammatory QTL 4 (*Nociq4*; MGI:5661503) (Figure 1A). This locus has not been previously detected in mouse genetic mapping studies. An approximate confidence interval for *Nociq4* was calculated using a 1-LOD drop from the peak SNP association, resulting in an interval width of 3.1 Mbp (proximal: rs246258668 [11.95 Mb]; distal: rs580950795 [15.07 Mb]). The maximum LOD (logarithm of odds) score for *Nociq4* is 5.71 and occurs over several SNPs, ranging from 14.26 – 14.33 Mb, giving a peak location of 14.29 Mb (GRCm38).

**Figure 1.**
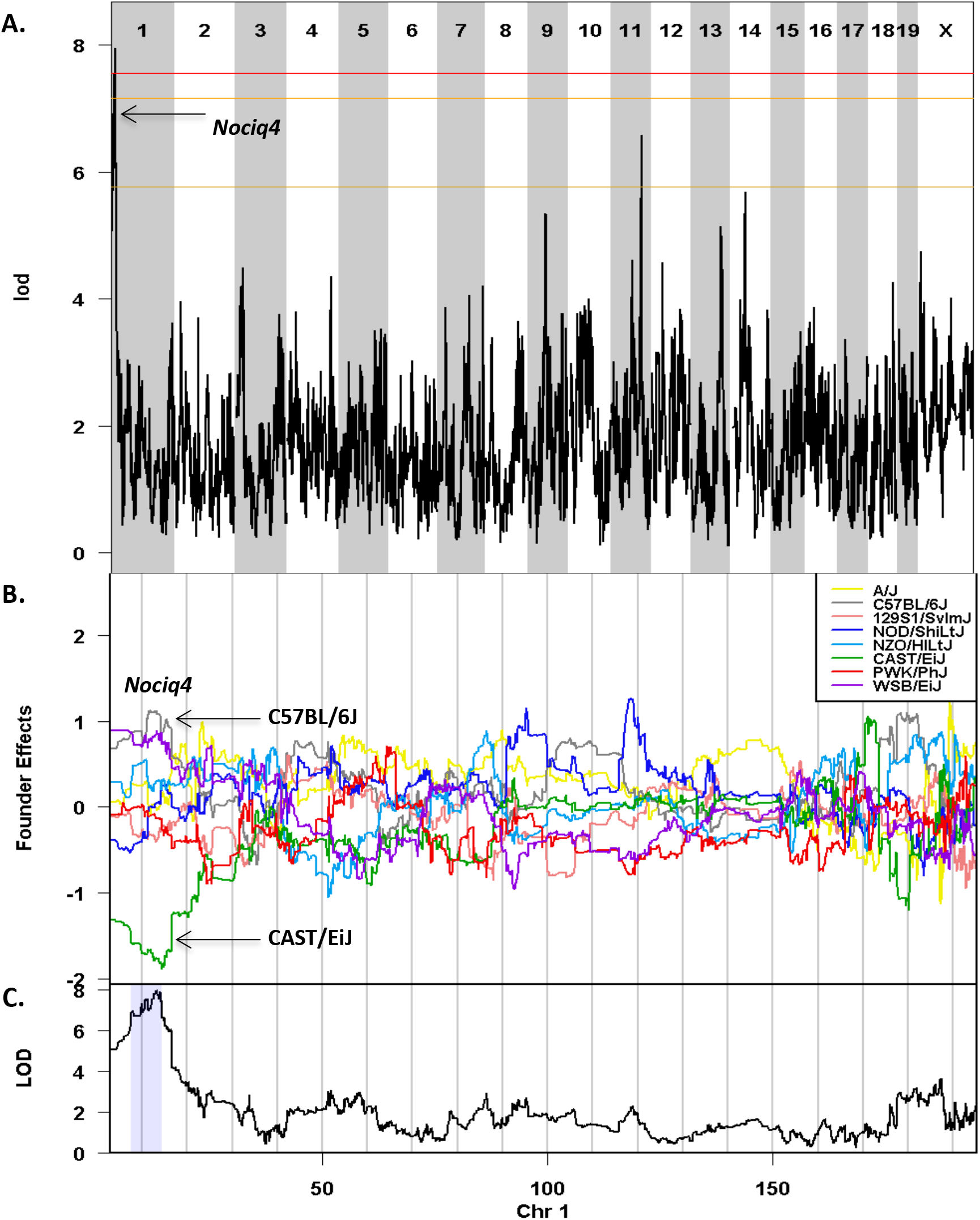
Late phase formalin response has a significant QTL on mouse chromosome 1 (*Nociq4*). **A.** Genome-wide scan for late phase response to formalin injection reveals a QTL with a peak LOD score of 5.71 at 14.25 Mb. Permutation-derived significance thresholds are marked by horizontal lines: 0.63 (bottom), 0.1 (middle), 0.05 (top). **B.** The founder allele effects show that the CAST/EiJ allele contributes to lower late phase formalin response sensitivity. Each line represents the effect of one of the eight founder alleles in DO mice. The differences between strains are significant when the LOD score in panel C is high. **C.** Genome scan for sensitivity to late phase formalin response on chromosome 1.

### Candidate gene analysis

The *Nociq4* locus contains putatively 43 candidate genes: 11 protein-coding, 20 non-coding RNA, and 12 unclassified (GRCm38; MGI Genes and Markers query performed May 2018, Feature Type “gene” [87]). Annotations obtained from Ensembl annotation version 75 [85] produced similar results (Figure 2C). Rigorous de novo genetic or experimental evaluation of each candidate gene is inefficient and costly, so we compiled existing functional, phenotypic, and expression annotations for each gene to identify candidates with high relevance to pain based on known experimental evidence (Supplemental Table S2).

**Figure 2.**
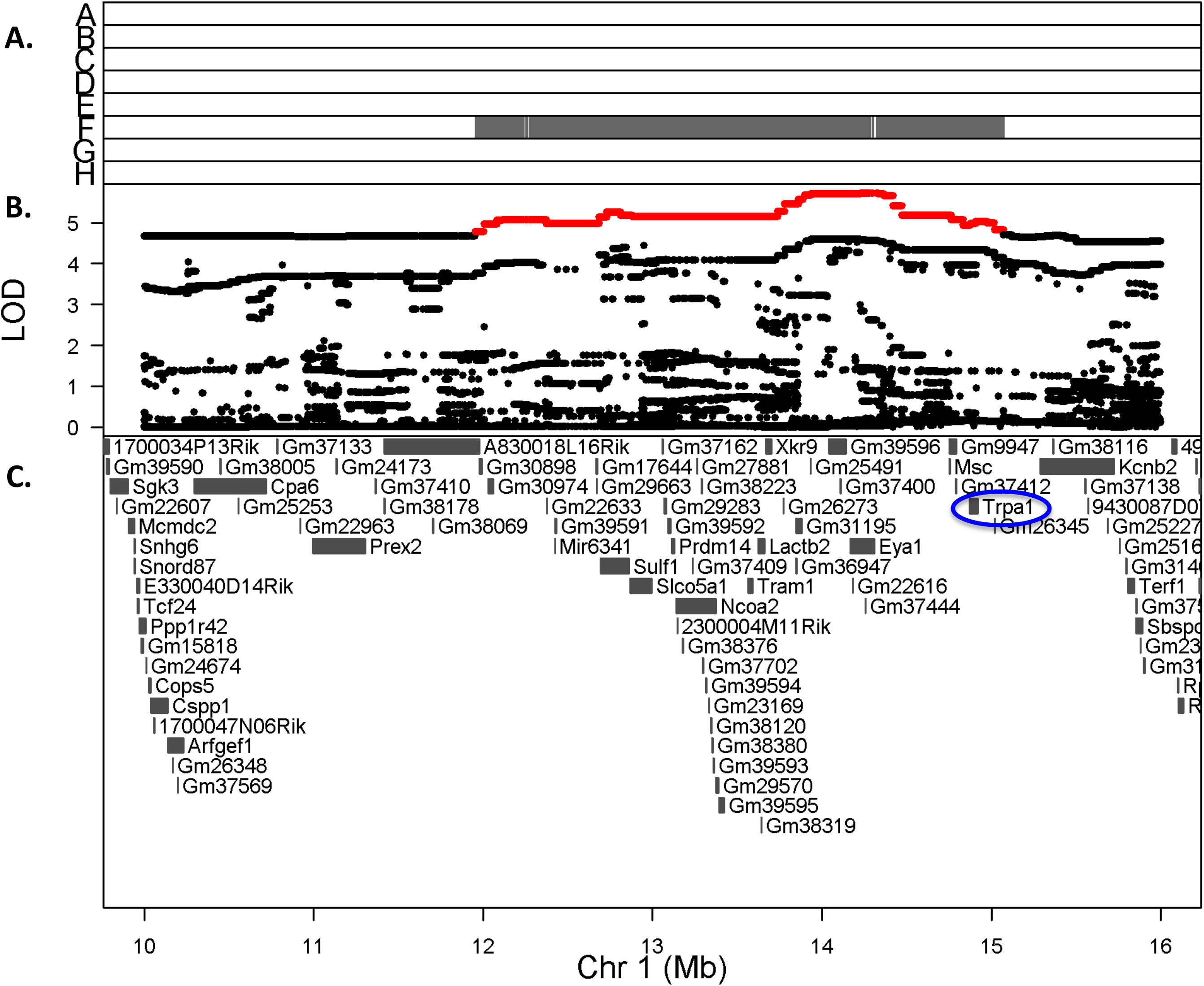
*Trpa1* lies within a genetically mapped region of chromosome 1 significantly correlated with late phase behavioral response to formalin injection. **A.** Minor allele frequency of the SNPs with the highest LOD score and shown in red in panel B. Strains: A, A/J; B, C57BL/6J; C, 129S1/SvImJ; D, NOD/ShiLtJ; E, NZO/HlLtJ; F, CAST/EiJ; G, PWK/PhJ; H, WSB/EiJ. SNPs for which only CAST/EiJ (F) contributes the alternate allele have the highest LOD scores at the *Nociq4* locus. **B.** LOD scores of SNP association mapping in the chromosome 1 QTL interval. Each point represents the LOD score from one SNP. Red SNPs represent a 1-LOD drop from the maximum LOD. **C.** Candidate protein coding genes underlying the *Nociq4* locus relative to mouse genome build GRCm38 and Ensembl annotation version 75 **[85]**. Top candidate gene *Trpa1* is circled in blue (Chr1:14.87-14.91 [-]).

At the time of this writing (June 2018), 16 of the 43 *Nociq4* candidate genes had no biological annotations or related functional data. All remaining candidates (27) had at least one annotated expression study reporting positive transcript identification in central nervous system (CNS), peripheral nervous system (PNS), or skeletal muscle tissue. Of these, only 6 had additional functional or phenotypic annotations related to nociceptive or other nervous system abnormalities: *A830018L16Rik, Prdm14, Ncoa2, Eya1, Msc,* and *Trpa1*. Annotations related to altered nociception and neuron responses point to *Trpa1* (transient receptor potential cation channel, subfamily A, member 1; MGI:3522699) as the most likely candidate gene in the region. The GeneWeaver GeneSet graph in Figure 3 corroborates these results, identifying *Trpa1* as the highest-ranking candidate gene in the *Nociq4* interval meeting both genetic and functional criteria [8].

**Figure 3.**
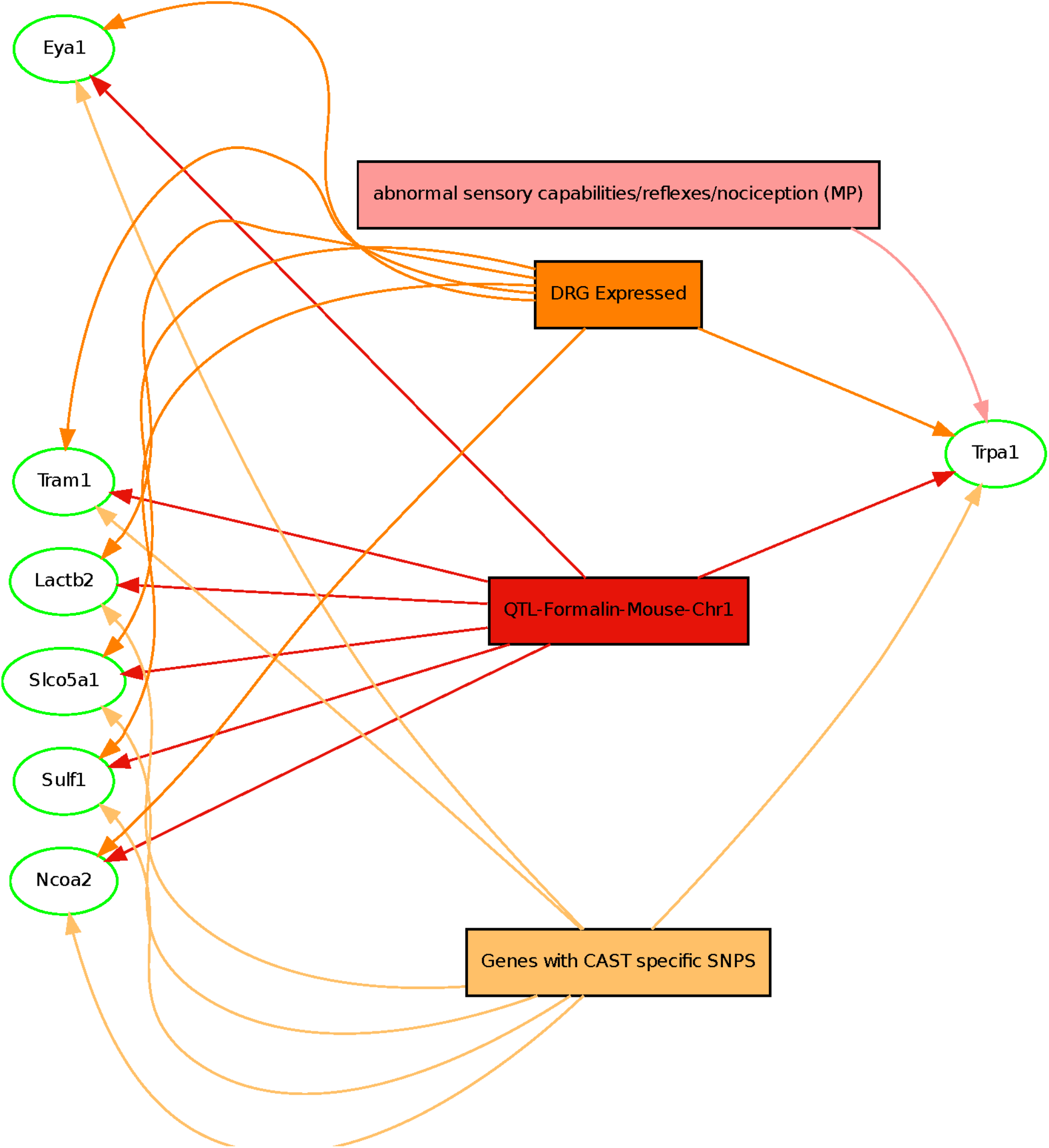
GeneWeaver analysis corroborates *Trpa1* as most likely *Nociq4* candidate gene. Output from the gene set graph tool showing *Trpa1* as the most highly connected gene. The genes are represented by oval shaped nodes, edges represent gene set membership, and the rectangular nodes represent gene sets retrieved from the GeneWeaver database. Other highly connected genes include *Eya1, Tram1, Lactb1, Slo5a1, Sulf1, Ncoa2*.

*Trpa1* is found in the plasma membranes of pain-detecting sensory nerves [64]. Gated by electrophilic compounds such as formalin, *Trpa1* is known to signal cell membrane deformation as well as noxious chemicals and temperatures [44]. Human TRPA1 is involved in inflammatory and neuropathic pain [57], and a point mutation at TRPA1 N855S is responsible for familial episodic pain syndrome [39]. Because of its capacity to respond to a wide variety of chemical compounds, *Trpa1* is considered critical for noxious chemical sensation, inflammatory signaling, and physiological and pathophysiological pain sensation [50]. It triggers beneficial avoidance behaviors and promotes longer-lasting biological responses such as inflammation, rendering it an attractive target for the treatment of both acute and chronic pain [18; 22].

### *CAST/EiJ alleles contribute to diminished late phase response at* Nociq4

Mice harboring CAST/EiJ alleles at *Nociq4* show a diminished late phase response to formalin (Figure 1B). Examination of the summed phenotypic scores of mice averaged over time within each haplotype carrier group (n=8) at the *Nociq4* peak (Chr1:14.25 Mb) reveals a unique behavioral response by mice harboring the CAST/EiJ allele at *Nociq4* – they do not appear to exhibit a late phase response to formalin injection despite an intact early phase response (Figure 4). Formalin injection typically induces a biphasic response in rodents, with behavioral plots showing two distinct peaks marking the onset, maintenance, and ending of acute and chronic pain states (0-10 mins and 10-60 mins, respectively). Most strains show the classic biphasic response curve, with early and late phase behavioral response peaks clearly visible (4A and 4C, respectively). During the early response phase (4A), mice exhibit an acute pain response regardless of DO founder haplotype at *Nociq4*. Mice harboring CAST/EiJ alleles at *Nociq4* do not appear to exhibit a late phase formalin response at 4C.

**Figure 4.**
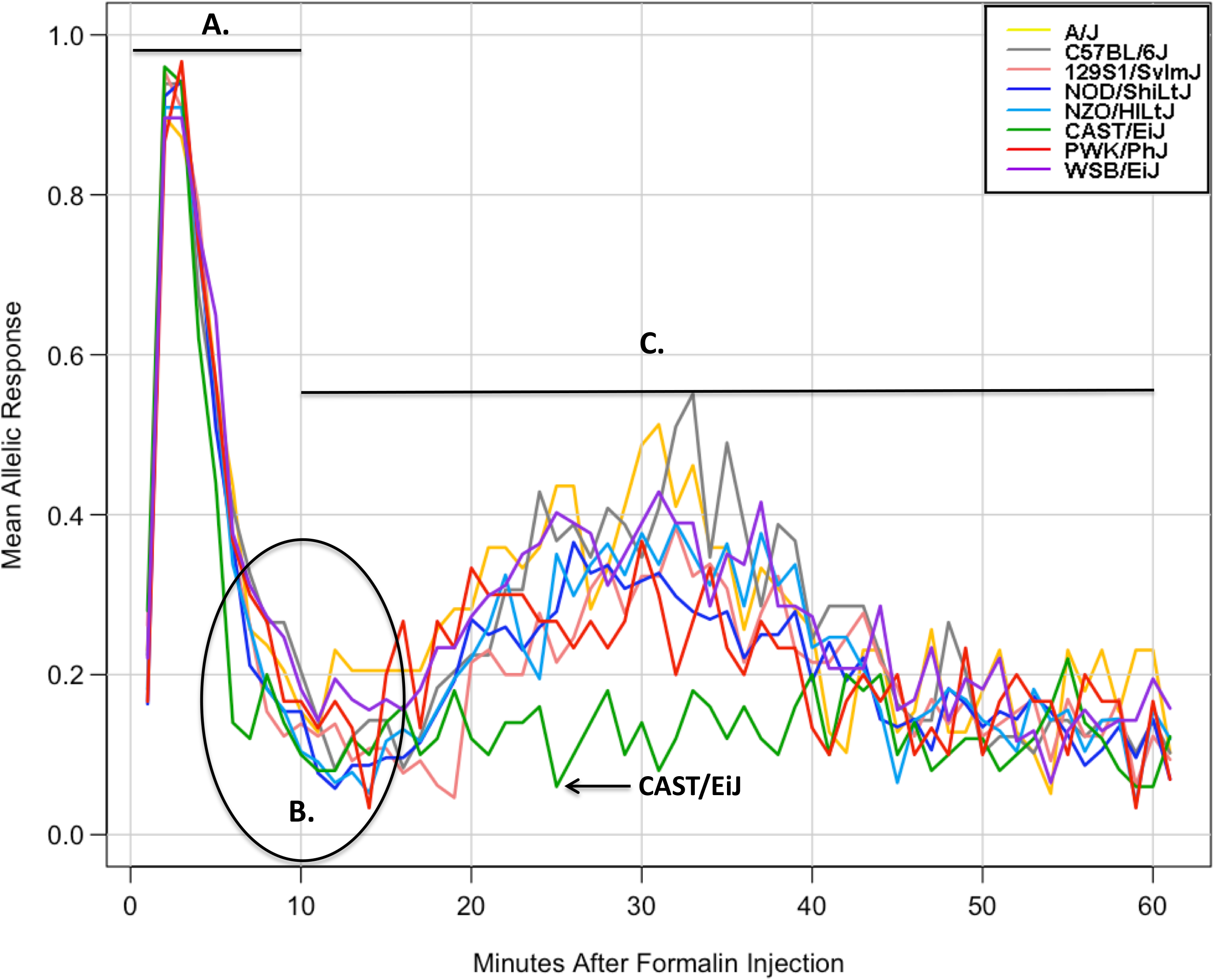
CAST/EiJ mice exhibit a strain-specific response to formalin injection at *Nociq4*. Averaging founder strain allele effects over time relative to *Nociq4* indicates a unique contribution by the CAST/EiJ allele to lower phenotypic response during late phase formalin injection. Each line represents the effect of one of the eight founder alleles in DO mice. **A.** All 8 DO founder strains respond similarly during the early (acute pain) phase following formalin injection (0-10 mins). **B.** All 8 strains exhibit a drop in phenotypic response ∼10 mins post-injection, which signifies the shift from the acute pain response (early phase) to the (late phase) chronic pain response. **C.** Only CAST/EiJ does not develop the “tonic” or chronic pain behavior typically associated with late phase behavioral response to formalin injection (10-60 mins).

### Trpa1 *SNP Analysis*

To more precisely identify the basis of the allelic affect pattern at *Nociq4* of a decreased late-phase response in mice harboring the CAST/EiJ allele at the locus, we used SNP data from NCBI dbSNP Build 150 [1; 69] and the Sanger Mouse Genomes Project version 5 (REL-1505) [37] to identify putative causal *Trpa1* variants unique to CAST/EiJ. Of the 201 SNPs identified, 190 are located in introns and have no known or predicted functional consequence (Supplemental Table S3). Eight of the remaining 11 SNPs are synonymous coding exon variants. The three remaining SNPs have consequences likely to influence the function or expression of *Trpa1*: missense variant rs32035600 (Val115Ile; exon 3), 3’ UTR variant rs215479411, and splice region variant rs239908314 (intron 21) (Table 1).

**Table 1.** *Trpa1* SNPs significant for late phase formalin response unique to CAST/EiJ. (Bold italics denote variants selected for characterization based on functional class).

| rsID | Position (bp)<br>(GRCm38) | Ref<br>Allele | Alt<br>Allele | *Trpa1* location <sup>a</sup> | Functional class <sup>b</sup> |
| --- | --- | --- | --- | --- | --- |
| ***rs215479411*** | ***14872864*** | ***C*** | ***A*** | ***3' UTR variant*** | ***3' UTR variant*** |
| rs240389458 | 14882175 | G | A | exon 22 | Synonymous variant |
| ***rs239908314*** | ***14884107*** | ***T*** | ***G*** | ***intron 21*** | ***Splice region variant*** |
| rs262385541 | 14884181 | C | T | exon 21 | Synonymous variant |
| rs245414067 | 14884229 | A | G | exon 21 | Synonymous variant |
| rs219630558 | 14898047 | G | A | exon 12 | Synonymous variant |
| rs233861326 | 14898248 | G | A | intron 11 | Splice region variant |
| rs243856490 | 14901865 | T | G | exon 8 | Synonymous variant |
| rs252121819 | 14904526 | C | T | exon 5 | Synonymous variant |
| rs248884581 | 14905986 | C | T | exon 4 | Synonymous variant |
| ***rs32035600*** | ***14910834*** | ***C*** | ***T*** | ***exon 3*** | ***Missense variant*** |
| rs238899246 | 14912367 | C | T | exon 2 | Synonymous variant |
| rs32036619 | 14912526 | G | A | intron 1 | Splice region variant |

<sup>a</sup> Source: NCBI dbSNP Build 150 [1; 69].<sup>b</sup> Source: Sanger Mouse Genomes Project version 5; REL-1505 [37].

Missense variant rs32035600 is located in the ankyrin (ANK) binding domain of *Trpa1* [30]. The *Trpa1* ANK binding domain is involved in the recognition of electrophilic compounds [33; 48] and calcium [25], which could imply variant effects on the channel’s sensing and gating functions. ANK binding domain mutations could also influence *Trpa1* regulation by modifying the channel’s ability to colocalize with its functional partner *Trpv1* [82]. 3’ UTR variant rs215479411 is most likely to affect *Trpa1* expression levels through mechanisms of post-transcriptional modification, such as binding and degradation by micro- or other non-coding RNAs [62]. The splice region variant (rs239908314) is located in *Trpa1* intron 21 and may regulate the expression of transcript isoforms *Trpa1*a (full-length) and *Trpa1*b (lacking exon 20). [86]. We functionally characterized missense variant rs32035600, 3’ UTR variant rs215479411, and splice region variant rs239908314 for causal impacts on *Trpa1* by measuring changes in channel electrophysiology, receptor binding affinity/colocalization with *Trpv1*, and transcript isoform expression levels. The workflow behind our functional experimental protocol is summarized in Figure 5.

**Figure 5.**
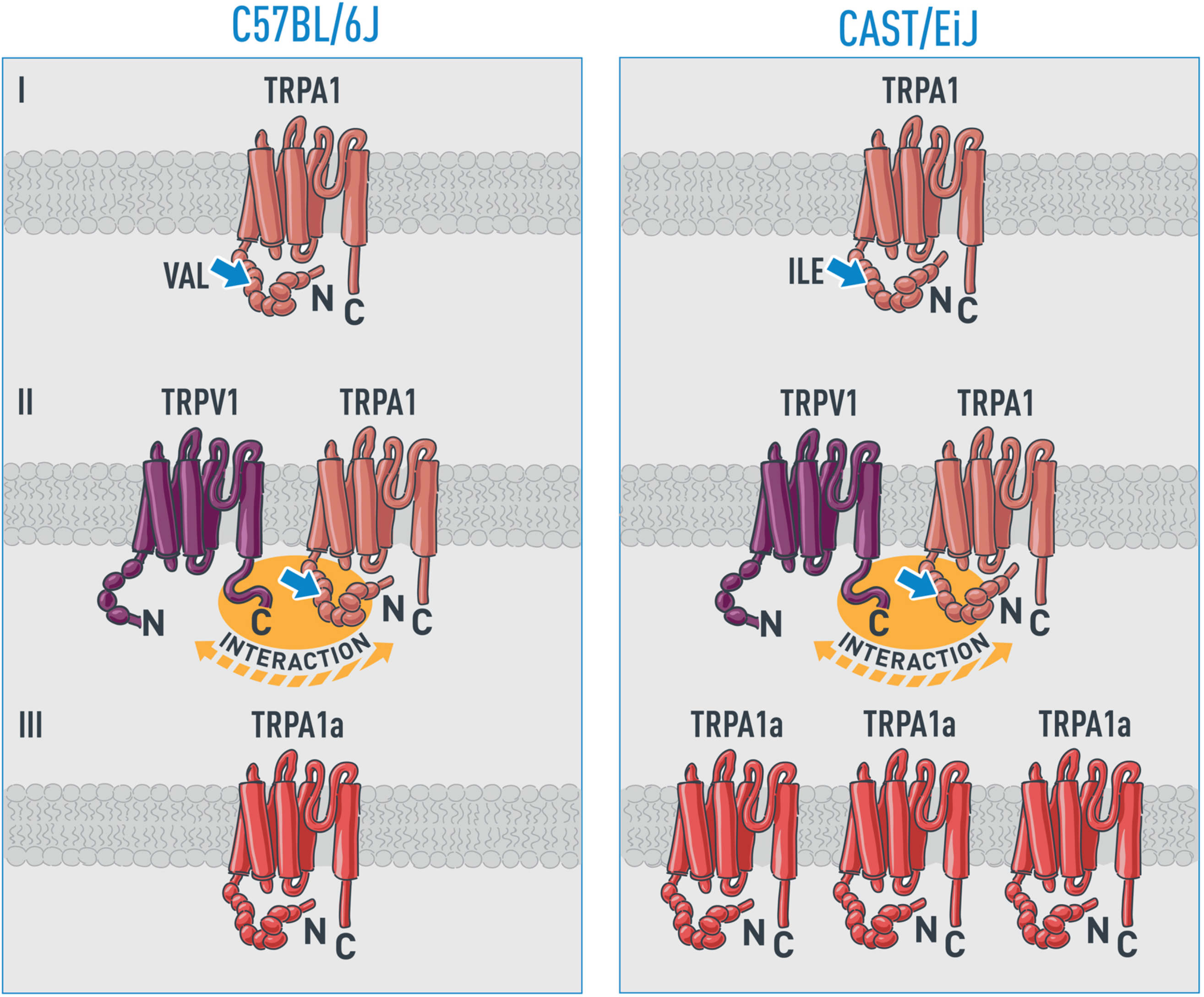
Schematic representation of potential functional effects of *Trpa1* candidate SNPs in C57BL/6J and CAST/EiJ. (I) SNP rs32035600 induces a valine 115 to isoleucine shift (a difference of one methyl group) between the strains in the ankyrin repeat domain of *Trpa1*. The slight side chain change may alter the structure and thus functional properties of the protein. (II) The same amino acid changing SNP, rs32035600, could affect *Trpa1* function in an allele specific manner by altering the *Trpv*1-*Trpa*1 interaction. (III) SNP rs239908314 in intron 21 regulates the expression and/or alternative splicing of the *Trpa1* transcript, affecting levels of *Trpa1*a and *Trpa1*b on the cell membrane. *Trpa1*a and *Trpa1*b expression levels could also be influenced by 3’ UTR SNP rs215479411 – if the same differential expression pattern is observed for both isoforms.

### Electrophysiology

Val115Ile encodes part of the *Trpa1* ankyrin (ANK) 2 binding domain (IPR002110), which is part of a larger chain of ANK repeats known as the ankyrin repeat domain (ARD). The *Trpa1* ARD facilitates cytoplasmic *Trpa1* inter-subunit interactions that may regulate channel assembly and/or facilitate conformational changes after co-factor binding or agonist-evoked gating [46]. Isoleucine is slightly more hydrophobic than Valine [52], suggesting rs32035600 may impact steric linking of the ANK repeat network structure. Our null hypothesis was that CAST/EiJ variant rs32035600 would have no effect on *Trpa1* channel gating and conductance compared to the C57BL/6J allele. We used whole-cell patch clamp recording to explore the electrophysiological consequence of CAST/EiJ rs32035600 on *Trpa1* function. HEK293T cells expressing the C57BL/6J and CAST/EiJ variants of *Trpa1* rs32035600 both showed robust responses to the application of mustard oil (MO) (Figure 6). There was no difference in current amplitude between C57BL/6J and CAST/EiJ *Trpa1*, or in the rise or decay time of MO-induced currents, suggesting that rs32035600 does not affect the gating and conductance of *Trpa1*. We note that the electrophysiological properties of the channel were investigated under a limited set of conditions, and therefore we cannot completely rule out an effect of the variant on channel properties.

**Figure 6.**
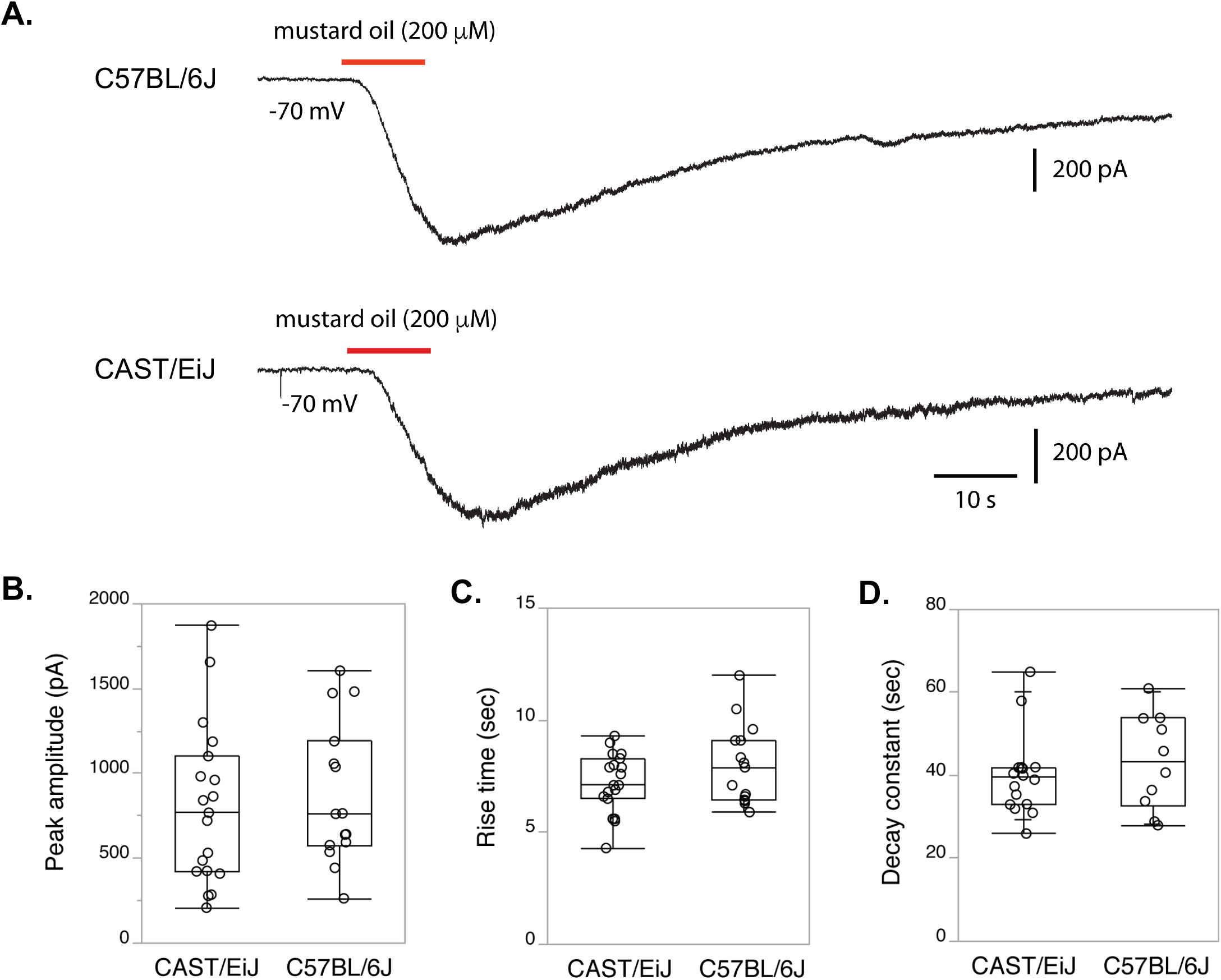
The *Trpa1* CAST/EiJ variant rs32035600 does not significantly alter *Trpa1* channel conductance in HEK293T cells. **A.** Both C57BL/6J and CAST/EiJ alleles of *Trpa1* variant rs32035600 showed robust responses to the application of mustard oil (MO) during whole-cell patch clamp recording in HEK293T cells 48-72h after transfection. **B.** There was no difference between C57BL/6J (n=15 cells) and CAST/EiJ (n=19 cells) *Trpa1* in current amplitude (*p* = 0.56). There was no difference between C57BL/6J and CAST/EiJ *Trpa1* in the rise time **(C.)** (10-90% of peak; *p* = 0.34) or decay time **(D.)** (*p* = 0.47) of MO-induced currents.

### *Co-Immuno-Precipitation of* Trpa1 *and* Trpv1

*Trpa1* and *Trpv1*, two ligand-gated non-selective cation channels, are known to be co-expressed in DRG [45], with a *Trpa1*-*Trpv1* interaction thought to be an important regulatory mechanism of persistent pain [82]. To determine if the amino acid changes resulting from the C57BL/6J / CAST/EiJ SNP (C:T; rs32035600) affect this interaction, protein extracts from the DRG of C57BL/6J and CAST/EiJ mice 30 minutes after formalin or saline injection were incubated with antibodies against *Trpv1*. The protein antibody complex was precipitated and subject to western blot analysis with antibodies against *Trpa1*. Quantification of blot intensity of *Trpa1*/*v1* ratios in both replicates revealed evidence of a sex by genotype effect such that the CAST/EiJ males have a heightened response to formalin that results in increased receptor co-IP and C57BL/6J females decrease receptor co-IP in response at 30 mins post formalin injection (Figure 7). Although interesting as a possible mechanism of sex x genotype interactions in pain sensitivity, this result does not explain the consistent effect of the CAST/EiJ allele on overall late phase response. Therefore, rs32035600 is unlikely to be responsible for variation in the formalin response through alteration of the *Trpa1*-*Trpv1* coupling.

**Figure 7.**
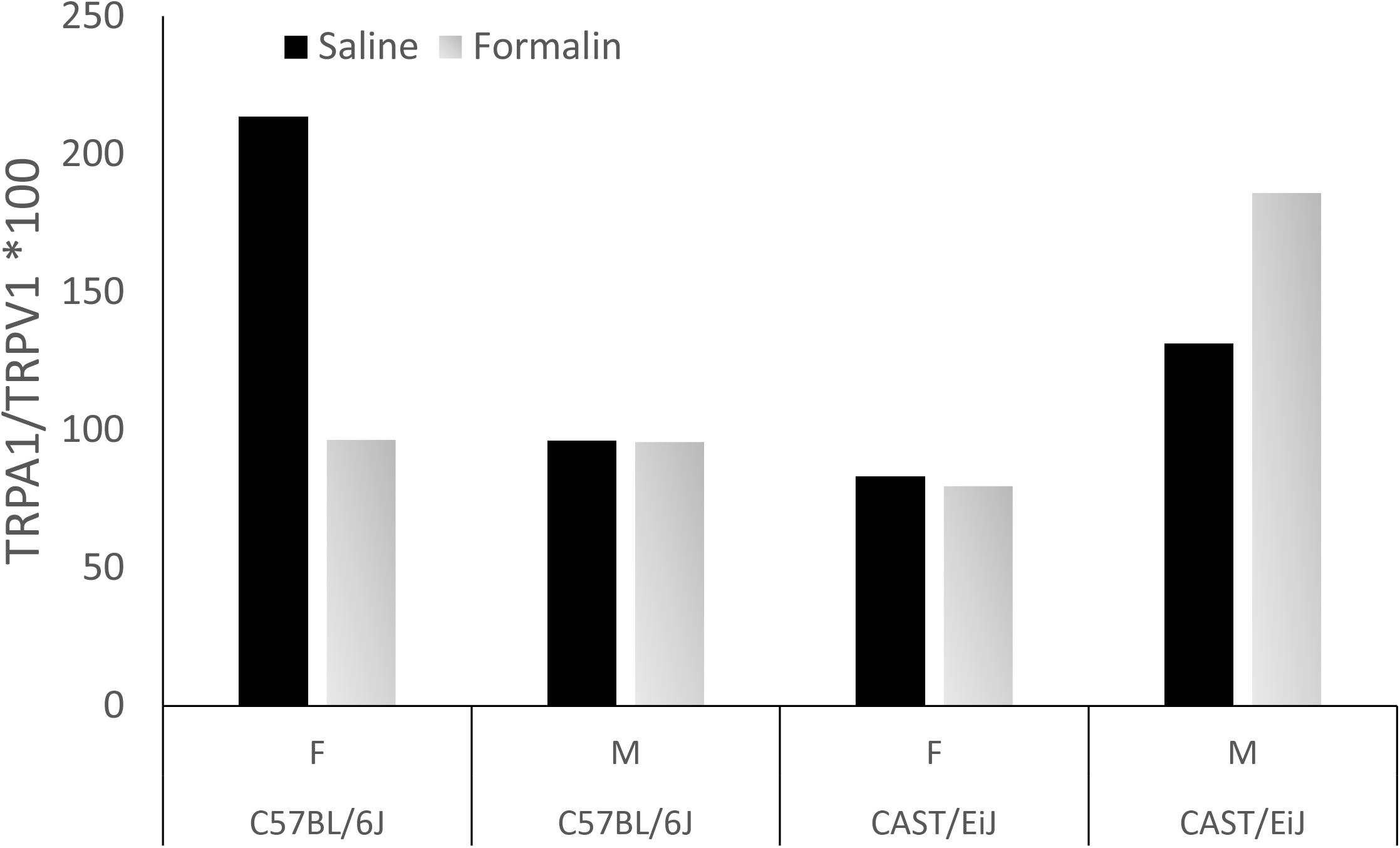
Quantification of western blot intensity of TRPA1-TRPV1 coupling shows a sex x genotype effect. CAST/EiJ males have a heightened response to formalin that results in increased receptor co-IP and C57BL/6J females decrease receptor co-IP in response at 30 mins post-formalin injection.

### Evaluation of expression regulatory variation

*Trpa1* has been shown to exist in mouse DRG in two isoforms: *Trpa1*a and *Trpa1*b. *Trpa1*a, the full-length transcript, is functionally conserved among mouse, rat, and human. The splice variant isoform, *Trpa1*b, lacks the transmembrane region encoded by exon 20 and appears to be non-functional as an ion channel. *Trpa1*b is known to physically interact with *Trpa1*a, enhancing the expression of *Trpa1*a on the cell’s plasma membrane. *Trpa1*b has been shown to regulate *Trpa1*a during the late stages of partial sciatic nerve ligation (PSL)-induced neuropathic pain and complete Freund’s adjuvant (CFA)-induced inflammatory pain [86].

The CAST/EiJ-specific SNP rs239908314 may regulate *Trpa1* alternative splicing by functioning as a splice region variant in *Trpa1* intron 21. We used qRT-PCR to explore the allelic effect of rs239908314 on *Trpa1* isoform abundance by quantifying *Trpa1*a and *Trpa1*b transcript levels in DRG from untreated CAST/EiJ (pain-resistant) and C57BL/6J (control) mice. Results show total *Trpa1* expression is approximately doubled in CAST/EiJ mice compared to age- and sex-matched C57BL/6J controls (Figure 8). When both *Trpa1*a and *Trpa1*b isoforms are considered, only *Trpa1*a is differentially expressed between the strains, expressed nearly three times higher in CAST/EiJ DRG. Our results indicate no difference in *Trpa1*b transcript abundance between CAST/EiJ versus C57BL/6J mice, which is consistent with work by Zhou et al. [86]. Our null hypothesis was that there was no difference in isoform expression levels in DRG from both untreated strains. We found instead a significant up-regulation of *Trpa1*a in CAST/EiJ mice, and a significant difference in isoform ratio between CAST/EiJ and C57BL/6J, with *Trpa1*a expressed nearly three times higher in CAST/EiJ DRG. If the differential abundance were a result of the 3’ UTR SNP (rs215479411, common to both *Trpa1*a and *Trpa1*b), the effect on post-transcriptional modifications would be expected to be the same in both isoforms, not just in the abundance of *Trpa1*a as observed. Our findings suggest a functional role for SNP rs239908314 in *Trpa1* isoform regulation, and a possible mechanism by which CAST/EiJ mice regulate their late phase response to formalin injection.

**Figure 8.**
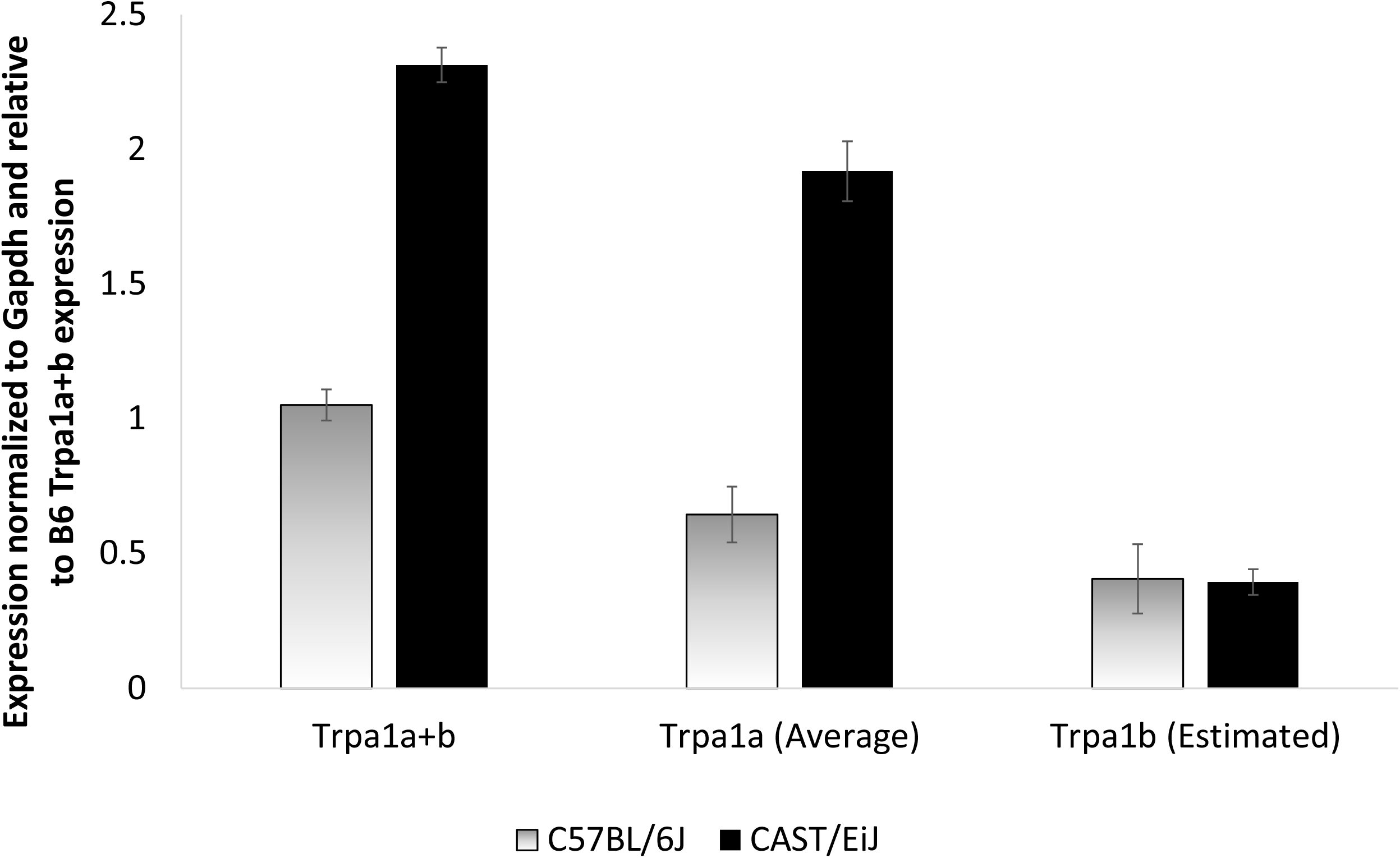
Quantitative-PCR shows higher average concentration of *Trpa1*a in the dorsal root ganglia (DRG) of naïve male and female CAST/EiJ compared to C57BL/6J mice. Total *Trpa1* expression is approximately doubled in CAST/EiJ mice compared to age- and sex-matched C57BL/6J controls. When both *Trpa1*a and *Trpa1*b isoforms are considered, only *Trpa1*a is differentially expressed between the strains, expressed nearly 3x higher in CAST/EiJ DRG.

## Discussion

Using genetic linkage mapping and genome wide SNP association mapping in a cohort of 275 DO mice, we identified a novel 3.1 Mbp late phase formalin response QTL, *Nociq4* (nociceptive sensitivity inflammatory QTL 4; MGI:5661503), on mouse chromosome 1 harboring 31 candidate genes. *Nociq4* harbors the well-known pain gene *Trpa1* (transient receptor potential cation channel, subfamily A, member 1), a cation channel governing acute and chronic pain in both humans and mice [4; 13; 38; 40; 41; 51].

We identified *Trpa1* as the most plausible candidate gene in the QTL region, noting a diminished late phase formalin response in mice harboring the CAST/EiJ allele at the locus. We characterized functional consequences of sequence variants in *Trpa1*: a missense variant resulting in a nonsynonymous amino acid change (rs32035600; Val115Ile) which could affect either electrophysiology or receptor colocalization, a 3’ UTR variant (rs215479411) which could affect overall transcript abundance, and a splice junction variant (rs239908314) which could affect transcript isoform expression. qRT-PCR analysis confirmed a three-fold expression difference in *Trpa1*a isoform abundance in untreated CAST/EiJ compared to C57BL/6J DRG, implicating *Trpa1* alternative splicing in diminished late phase formalin response.

Experimental evidence in rodents has shown that tonic (persistent) pain, similar to the chronic pain experienced by humans, is modulated by different CNS mechanisms than acute pain [79]. The first phase of the formalin test (0-10 minutes post-injection) is caused by intense neuronal activity in the spinal cord and serves as a model of acute pain [59; 79]. The second behavioral phase (occurring 10-60 minutes post-injection) is mediated by sensitization of spinal cord nociceptors and serves as a model of human chronic pain [59]. Spinal cord levels of c-fos, substance P, and excitatory amino acids also increase after formalin injection [79], inducing central sensitization via an excited nociceptive state similar to that observed in human chronic pain conditions [73; 74]. In the present study, we report that a point mutation in *Trpa1* (rs239908314) significantly reduces or even eliminates DO behavioral response during the late phase of the formalin test. This observation suggests that CNS sensitization mechanisms are critical for advancing the shift from acute to chronic pain and lends support to the idea that formalin induces a tonic pain state via CNS sensitization.

*Trpa1* is regulated by epigenetic modifications as well as non-coding RNAs [13; 31; 62; 75]. Candidate gene analysis predicted the presence of 20 non-coding RNAs within the *Nociq4* region, including two microRNAs (miRNAs). Functional, phenotypic, and expression annotations for these genes are currently incomplete, and therefore, potential interactions between them and *Trpa1* or other *Nociq4* candidates may be missed. We identified 3’ UTR variant rs215479411 as a possible *Trpa1* functional variant contributing to decreased late phase formalin response at *Nociq4*. We hypothesized a role for rs215479411 in *Trpa1* expression regulation based on the ability of miRNAs to degrade target transcripts by adhering to specific 3’ UTR binding sites [62]. Our qRT-PCR data suggest that rs215479411 does not alter the post-transcriptional abundance of *Trpa1*, however, as only transcriptional isoform *Trpa1*a was found to be differentially expressed between CAST/EiJ and C57BL/6J.

The interaction of some of these expression regulatory mechanisms with sex hormones or developmental sex differences in a genotype-specific manner could account for the complex pattern of findings we obtained in our analysis of *Trpa1* and *Trpv1* clustering. Many other regulatory mechanisms are possible and this finding could merit further confirmation and investigation. Genetic variation has been previously shown to influence both the magnitude and direction of sex differences in acute thermal nociception, and this same complexity no doubt exists for chronic pain [53]. Expression-QTL (eQTL) studies of *Trpa1* and other pain-related genes are warranted to gain new insights into the molecular genetic networks governing gene expression during acute, chronic, and pain-free states.

*Trpa1* is a polymodal chemosensor expressed primarily in nociceptive neurons of peripheral ganglia. It acts as a high-threshold chemo- and mechanosensor that integrates painful mechanical stimuli with other noxious signals [82]. Human TRPA1 is of particular interest as a drug target because of its expression in nociceptor sensory neurons and its capacity to transduce a wide variety of noxious chemical stimuli into action potentials [24]. Pharmaceutical TRPA1 antagonists developed to date have proven most useful as *in vivo* and *in vitro* tools for studying TRPA1 biology [18]. This is perhaps due to the preferential activation of the channel by exogenous electrophilic agonists [9; 11; 12; 14; 35; 49]. In order for TRPA1 to be regarded as a suitable target for pain and other disorders, it must be active in the context of a pathological state [18]. TRPA1 has been found to play an important role in linking the presence of oxidative stress to inflammatory and neuropathic pain through the role of endogenous agonists such as oxidized lipids [80] and H_2_O_2_ [5]. Because spinal activation of TRPA1 can be either nociceptive or antinociceptive [18], both antagonists and agonists of TRPA1 may have utility for pain relief.

TRPA1 activity undergoes functional desensitization through multiple cellular pathways which are not yet fully understood [3; 36]. Agonist exposure can increase the level of *Trpa1* expressed on the cell membrane surface, suggesting a putative mechanism by which alternative splice variant rs215479411 modulates decreased late phase response to formalin injection in CAST/EiJ mice. SNP rs215479411 may modulate *Trpa1* agonist response by lending to increased cellular membrane expression of *Trpa1*a in CAST/EiJ, leading to quicker functional desensitization of the receptor compared to other DO inbred founder strains. This hypothesis is supported by Zhou and colleagues [86], who report dynamic changes in *Trpa1*a and *Trpa1*b expression levels during inflammatory and neuropathic pain conditions. Further investigation of this functional effect on *Trpa1* activity in humans is warranted to determine the potential clinical utility of the mechanism.

The work described in this article represents the first application of DO mice to chronic pain genetics research. Taken together, our results demonstrate that high-precision mapping of pain-related genetic variants can be achieved with moderate numbers of DO animals, representing a significant advance in our ability to leverage the mouse as a tool for the discovery of pain-related genes and therapeutic targets. Precise genetic analysis enabled us to identify not just the target gene, but three putative mechanisms of genetic effects on the phenotype. *Trpa1*a/b isoform regulation is involved in sparing of the intact acute pain response, which is a necessary sensory function, while specifically blocking the late phase response. Our results suggest that facilitating the effects of the *Trpa1*a isoform may have beneficial and specific effects on chronic but not acute pain. Applying our method of discovery to other pain-related traits may implicate other pain-relevant genes and novel variant contributions to pain response, facilitating the informed identification of therapeutics aided by the use of genetic precision to prioritize specific sub-molecular targets.

## Acknowledgements

Dr. Recla reports a grant from the US Department of Defense (DOD) and grants from the National Institutes of Health (NIH) during the conduct of the study: This work was funded in part by DOD grant W81XWH-11-1-0762 (CJB). This project was also supported by NIH R01 DA 37927 and NIH R01 AA 18776 to EJC. We gratefully acknowledge The Jackson Laboratory (JAX) Scientific Services supported by NIH P30 CA034196, as well as the technical efforts of Andrew Garrett and Robert Burgess (JAX) for their work on HEK cell transfections. The authors declare no other conflicts of interest.

